# Impact of sex and glial tau expression on heat shock protein induction in a *Drosophila* model of tauopathy

**DOI:** 10.1101/2025.07.10.664197

**Authors:** Margot Whitmore, Maeve Coughlan, Martha A. Kahlson, Jaasiel Alvarez, Louisa Zebrowski, Kenneth J. Colodner, Kathryn A. McMenimen

**Author notes:** **Corresponding authors:** Kenneth J. Colodner and Kathryn A. McMenimen.

## Abstract

Heat shock proteins (Hsps) are central components of the cellular stress response and serve as the first line of defense against protein misfolding and aggregation. Disruption of this proteostasis network is a hallmark of neurodegenerative diseases, including tauopathies – a class of neurodegenerative diseases characterized by intracellular tau accumulation in neuronal and glial cells. Although specific Hsps are enriched in glial cells, and some have been shown to directly bind tau and influence its aggregation, the broader interplay between Hsps and tau remains poorly understood. In particular, it is unclear whether tau expression affects the heat shock response, and whether this interaction is modulated in a sex-specific fashion. Here, we used a *Drosophila* model of tauopathy to examine both inducible and constitutive Hsp expression in response to heat stress in the context of glial tau expression. We found that Hsp expression displays sexually dimorphic expression patterns at basal levels and in response to heat stress. Moreover, tau expression in glia disrupts the normal induction of specific heat shock proteins following heat stress. This work provides new insight into how tau interacts with the cellular stress response, and highlights sex-specific differences in Hsp regulation. Understanding these molecular connections is crucial to understanding how the presence of tau in glial cells influences the stress response, and potentially contributes to tauopathy pathogenesis.

## Introduction

Neurodegenerative diseases are primarily characterized by the accumulation of misfolded protein aggregates. A critical line of defense against producing these aggregates is the heat shock response, which activates a global network of molecular chaperones(1, 2). Heat shock proteins (Hsps), originally discovered in *Drosophila* and named for their critical response to heat stress (3–5), are key components of this network. These chaperones maintain essential processes including: protein transport (6), non-native protein degradation, protein complex assembly(7, 8), protein metabolism(9, 10), and protein aggregate disaggregation. (11) Disruptions in heat shock protein expression and function, therefore, have been implicated in several neurodegenerative disorders.(12–15)

Tauopathies are a broad class of neurodegenerative diseases defined by the pathological intracellular accumulation of the microtubule-associated protein, tau. These diseases, which include Alzheimer’s disease (AD), feature tau aggregation in both neuronal and glial cells. (16) Glial tau pathology is often found in brain regions with neuronal tau pathology (17, 18), and while its exact role in disease progression remains unclear, it can be used as a diagnostic hallmark in specific tauopathies. (19, 20) Animal models have demonstrated that glial tau expression can impair glial cell function (21–23), yet the molecular mechanisms linking tau accumulation and glial dysfunction remain incompletely understood.

Glial cells express Hsps, and it is not yet clear whether the formation of fibrillar glial tau inclusions reflects a failure of the Hsp network. Several studies suggest that Hsps are involved in regulating tau function and turnover, particularly in preventing the aggregation of pathologically modified tau. (24, 25) In response to proteasomal stress, chaperones, such as Hsp27, Hsp70/Hsp40/Hsp110, and Hsp90, are recruited to the cytoskeleton, where they can interact with tau. (24, 26–28) Moreover, Hsp27 and Hsp70 are robustly expressed in glial cells and have been observed to localize with tau inclusions in glia (26, 29, 30). Astrocytic Hsp27 is particularly upregulated in brain regions where neuronal tau tangles are prominent (31, 32), and may even be secreted from astrocytes to modulate inflammation and tau aggregation in neighboring neurons (33). These findings suggest that glial Hsps could influence tauopathy disease progression, but the relationship between glial tau accumulation and Hsp expression remains unclear.

Hsps are synthesized at elevated levels in response to cellular stress, and emerging evidence across organisms and tissues suggests that Hsps are differentially regulated between sexes. For example, in rodents, females display elevated levels of Hsp70, and decreased levels of Hsp27 and Hsp90, in heart tissue compared to males (34). In *Drosophila*, female flies exhibit greater thermotolerance than males, though the extent to which differential Hsp expression underlies this observation remains to be determined (35). Notably, small Hsp expression in *Drosophila* is developmentally regulated in a sex-specific and tissue-specific fashion (36), but our understanding of sex-specific Hsp regulation remains incomplete.

While sex-specific effects on the incidence of tauopathies have been described (37–40), and Hsp dysregulation has been implicated in aging and tauopathies (41–43), the intersection of sex and tau expression on Hsp dynamics remains poorly defined. To address this gap, we examined the expression of eight Hsps in response to early glial tau expression and heat stress in both male and female *Drosophila*. Our goal was to uncover potential sex-specific regulation of the heat shock response in the context of both physiological and disease-related stress.

## Results

### Basal expression of Heat shock proteins (Hsps) reveals sex- and Hsp-specific differences in day 10 control and glial tau-expressing flies

To assess basal (no heat stress) expression levels of Hsps and examine sex-specific differences, we performed qPCR gene expression analysis of eight Hsp genes in day 10 male and female control flies (Figure 1A). We used driver control flies (genotype: *repo-GAL4, tubulin-GAL80^TS^/+*) for these analyses to allow for a direct comparison to the genetic background of glial tau transgenic flies (genotype: *repo-GAL4, tubulin-GAL80^TS^, UAS-Tau^WT^/+*) previously established as a *Drosophila* model of glial tauopathy. (21, 44) In control flies, females exhibited significantly reduced basal expression of several inducible Hsp genes - *Hsp23*, *Hsp26*, and *Hsp70*, compared to males. In contrast, *Hsp27* expression was uniquely elevated in females relative to control males (Figure 1A). The constitutively expressed Hsps (*Hsc70* and *Hsp60*) showed no significant sex-dependent differences (Figure 1A).

**Figure 1.**
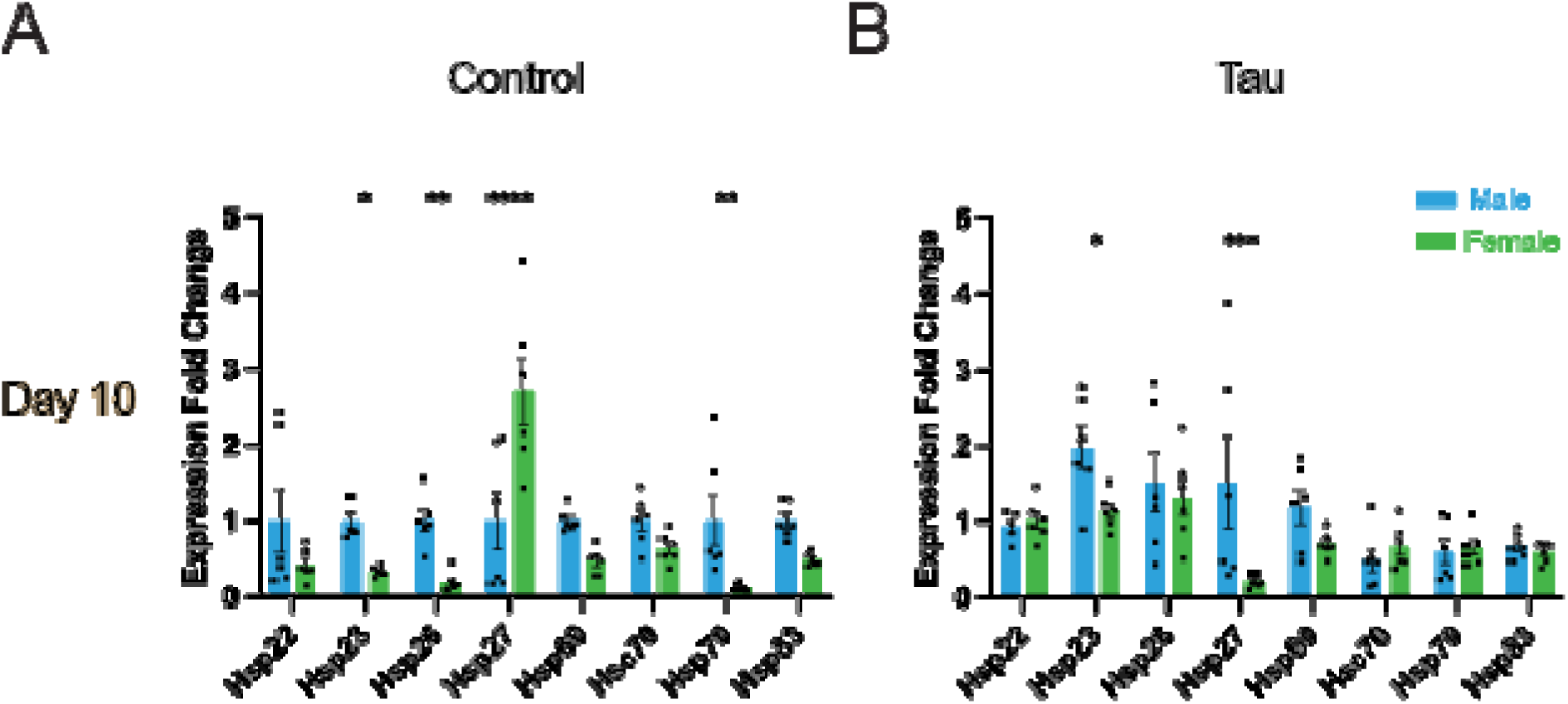
Basal male and female Hsp expression levels in control and tau transgenic flies at day 10. (A-B) Basal mRNA expression levels of six inducible (*Hsp22*, *Hsp23*, *Hsp26*, *Hsp27*, *Hsp70*, and *Hsp83*) and two constitutive (*Hsp60* and *Hsc70*) heat shock proteins in male and female control and tau transgenic flies. (A) Day 10 control flies display differential expression between male and female flies for specific Hsps. (B) Day 10 tau transgenic flies display differential expression between male and female flies for specific Hsps. Female data for both control and tau flies are normalized to day 10 male control flies for each Hsp. Data are presented as mean + SEM (n = 6) and analyzed using two-way ANOVA with Tukey’s multiple comparisons, * p < 0.05; ** p < 0.01; *** p < 0.001; **** P < 0.0001.

We next examined basal Hsp expression levels in glial tau transgenic flies at day 10 (Figure 1B). We focused our analysis on day 10 flies to capture early effects of glial tau expression, prior to the onset of significant tau pathology and neurodegeneration occurs in this model as previously described. (21) Similar to control flies, we found that day 10 male tau transgenic flies showed significantly higher expression of *Hsp23* relative to female day 10 tau transgenic flies (Figure 1B). However, in contrast to the control condition, *Hsp27* was elevated in male tau transgenic flies compared to females, reversing the sex-specific pattern observed in controls (Figure 1A, B). The ATP-dependent chaperone, *Hsp70*, also exhibited sexual dimorphic expression in control flies, with higher levels in male flies than females (Figure 1A). This elevated Hsp70 expression was not seen in tau transgenic flies, suggesting that glial tau expression disrupts normal regulation of *Hsp70* expression (Figure 1A, B). Expression of the two constitutively expressed chaperones, *Hsp60* and *Hsc70*, remained stable across sexes and showed no significant changes in response to glial tau expression (Figure 1A, B). In summary, basal heat shock protein expression reveals a trend of reduced expression in females relative to males, with *Hsp27* as a notable exception in control flies (Figure 1A). This trend for reduced expression of select Hsps in females is less pronounced, but still present, in glial tau transgenic flies (Figure 1B).

### Glial tau expression and heat stress differentially influence small heat shock protein expression in male and female flies

To determine the effect of heat stress and glial tau expression on sHsp expression, we examined expression levels for four small heat shock proteins (*Hsp22*, *Hsp23*, *Hsp26*, and *Hsp27*) in male and female control and glial tau transgenic flies (Figure 2). Expression levels were analyzed in the presence and absence of heat stress, and are presented as fold changes relative to basal male control flies. As expected, all four sHsps were significantly upregulated in response to heat stress across genotypes and sexes. *Hsp22*, a mitochondrial sHsp, exhibited robust heat-induced expression in both male and female flies (Figure 2A, B), and glial tau expression did not significantly alter *Hsp22* induction in either sex. *Hsp23* also showed strong induction following heat stress in both sexes (Figure 2C, D). However, glial tau expression significantly reduced *Hsp23* induction in males, while females were unaffected by glial tau expression (Figure 2C, D). A similar pattern was observed for *Hsp26*. Both female and male flies exhibited robust heat-induced expression, while glial tau expression significantly decreased *Hsp26* induction in males and not females (Figure 2E, F). *Hsp27* expression increased in response to heat stress in both males and females, and was the only sHsp where the presence of glial tau expression augmented the heat stress-induced increase in *Hsp27* in females only (Figure 2G, H).

**Figure 2.**
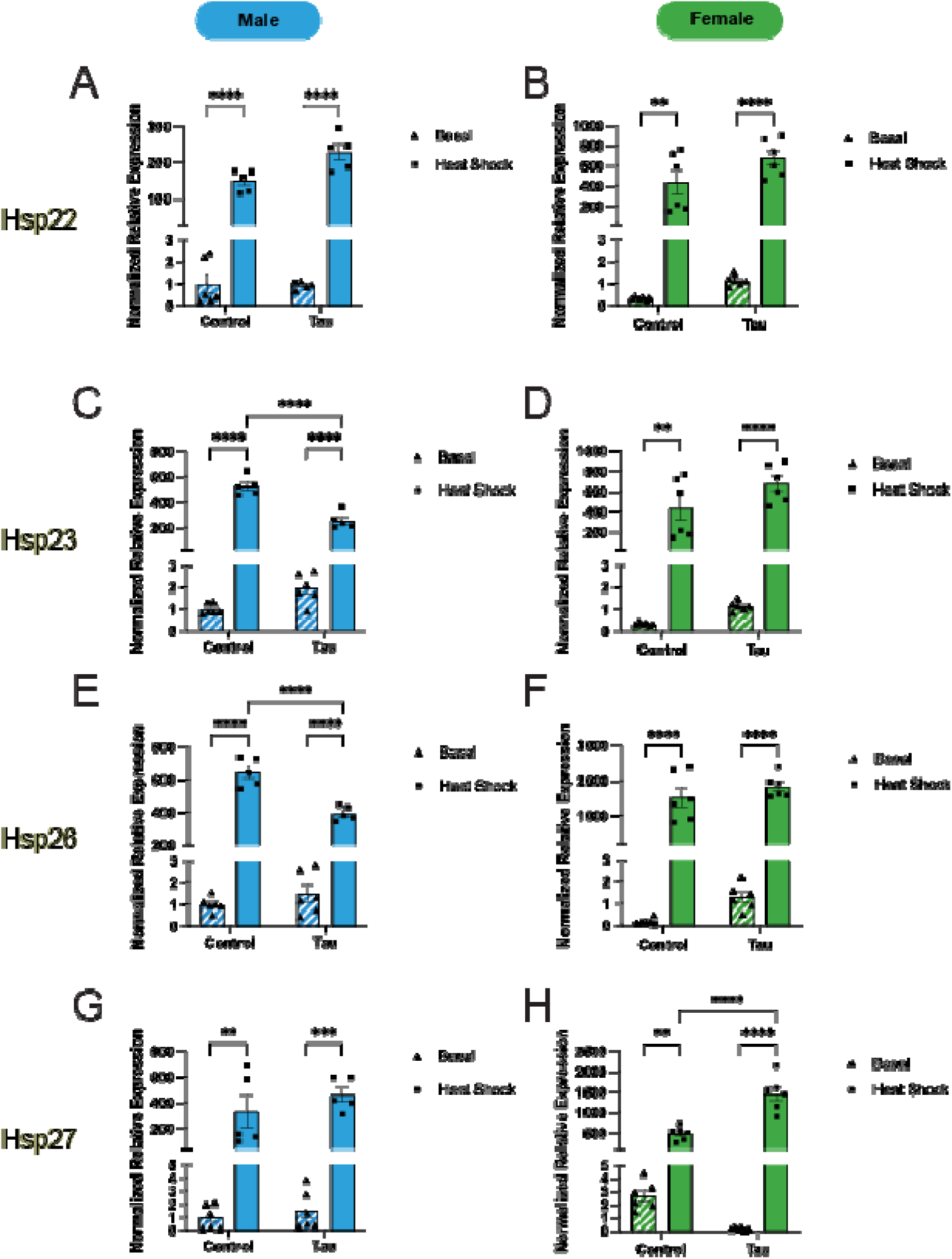
Small Hsp (sHsps) expression varies in male and female flies in response to glial tau expression and heat stress. (A-H) Relative mRNA expression levels of sHsps (*Hsp22*, *Hsp23*, *Hsp26*, *Hsp27*) for male (A, C, E, G) and female (B, D, F, H) flies under basal and heat stress conditions, comparing control and glial tau transgenic flies. Expression values are normalized to day 10 basal male control flies for each Hsp. Data are presented as mean + SEM (n = 6) and analyzed using two-way ANOVA with Tukey’s multiple comparisons * p < 0.05; ** p < 0.01; *** p < 0.001.

In summary, while heat stress reliably induces the upregulation of all tested sHsps, glial tau expression modulates this response in a gene- and sex-specific manner. Notably, tau suppresses *Hsp23* and *Hsp26* induction in male flies, and enhances *Hsp27* induction in female flies.

### Heat stress and glial tau expression differentially affect *Hsp70* and *Hsp83* expression in male and female flies

To assess the combined effects of heat stress and glial tau expression on the expression of large Hsps, we analyzed *Hsp70* and *Hsp83* transcript levels in male and female flies under all experimental conditions. Again, expression levels are presented as fold changes relative to male basal control flies (Figure 3). As expected, *Hsp70* was strongly induced by heat stress in both sexes (Figure 3A, B). However, the magnitude of induction was markedly higher in females (>2000-fold) than in males (300-fold). Notably, in females, glial tau expression further enhanced heat-induced Hsp70 expression, while this effect was not observed in males. *Hsp83* expression exhibited a similar pattern, though the fold change in response to heat stress for both sexes (<20-fold) was smaller (Figure 3C, D). In males, glial tau expression did not significantly alter *Hsp83* induction in response to heat stress (Figure 3C). In contrast, females exhibited a significant increase in *Hsp83* expression in response to heat stress when glial tau was present (Figure 3D).

**Figure 3.**
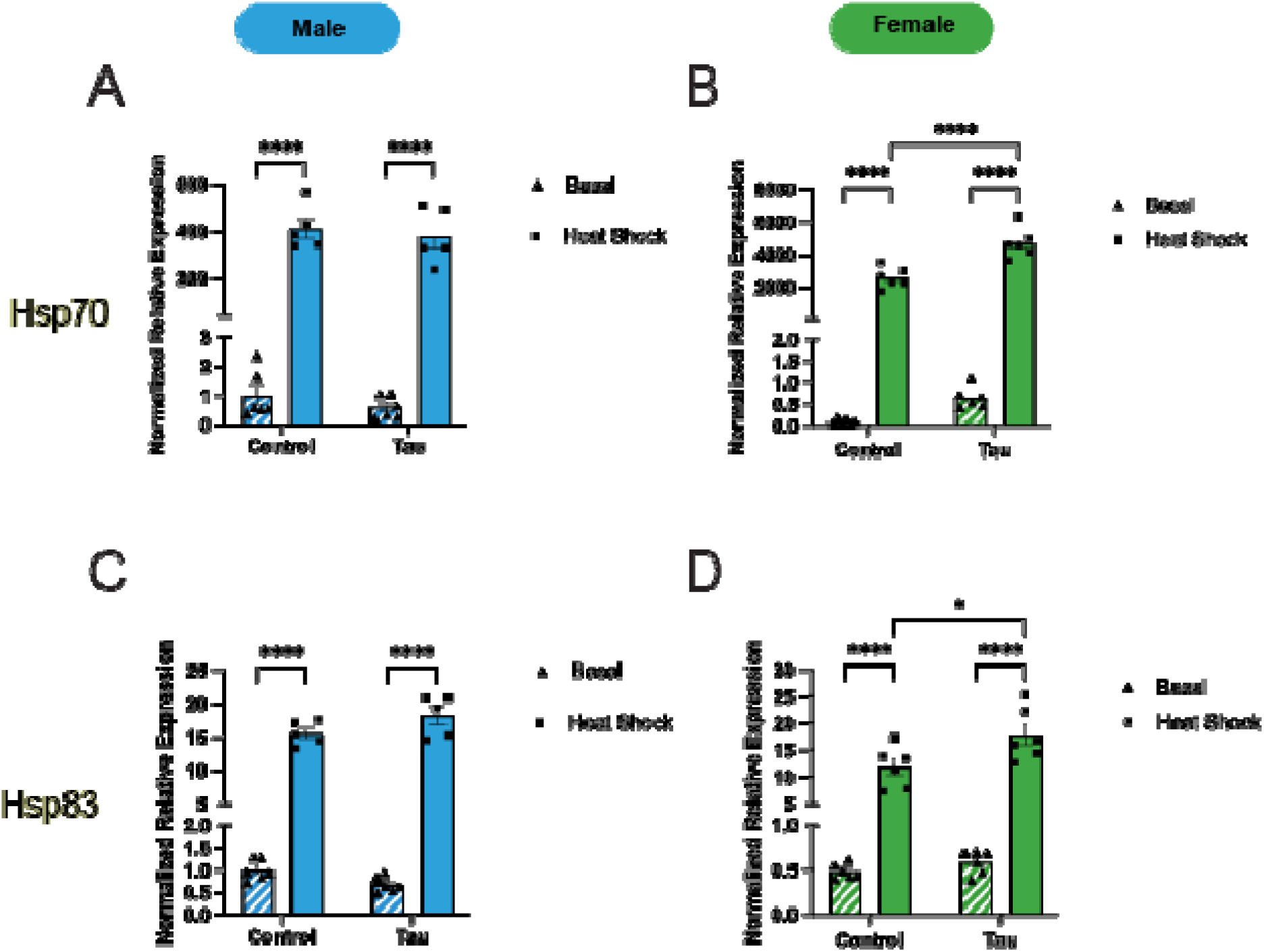
Large Hsp expression varies in day 10 male and female flies in response to glial tau expression and heat stress. (A-D) Relative mRNA expression levels of two large inducible heat shock proteins, *Hsp70* (A-B) and *Hsp83* (C-D), for male and female flies under basal and heat stress conditions, comparing control and glial tau transgenic flies. Expression values are normalized to day 10 basal male control flies for each Hsp. Data are presented as mean + SEM (n = 6) and analyzed using two-way ANOVA with Tukey’s multiple comparisons, * p < 0.05; ** p < 0.01; *** p < 0.001, **** P < 0.0001.

Together, these results show that *Hsp70* and *Hsp83* are differentially regulated by glial tau expression and heat stress in a sex-dependent fashion, with females showing greater responsiveness to combined heat stress and glial tau expression.

### Constitutive Hsps exhibit sexually dimorphic responses to heat stress and glial tau expression

To assess how constitutively expressed Hsps respond to heat stress and glial tau expression, we examined expression levels of *Hsp60* and *Hsc70*, reporting changes normalized to male basal control flies. *Hsp60* expression remained unchanged in response to heat stress and glial tau expression in male flies (Figure 4A). In contrast, female flies exhibited a significant induction of *Hsp60* only when both heat stress and glial tau expression were present (Figure 4B). Neither heat stress nor the presence of glial tau expression alone was sufficient to alter *Hsp60* expression in females (Figure 4B), suggesting a combinatorial effect specific to females.

**Figure 4.**
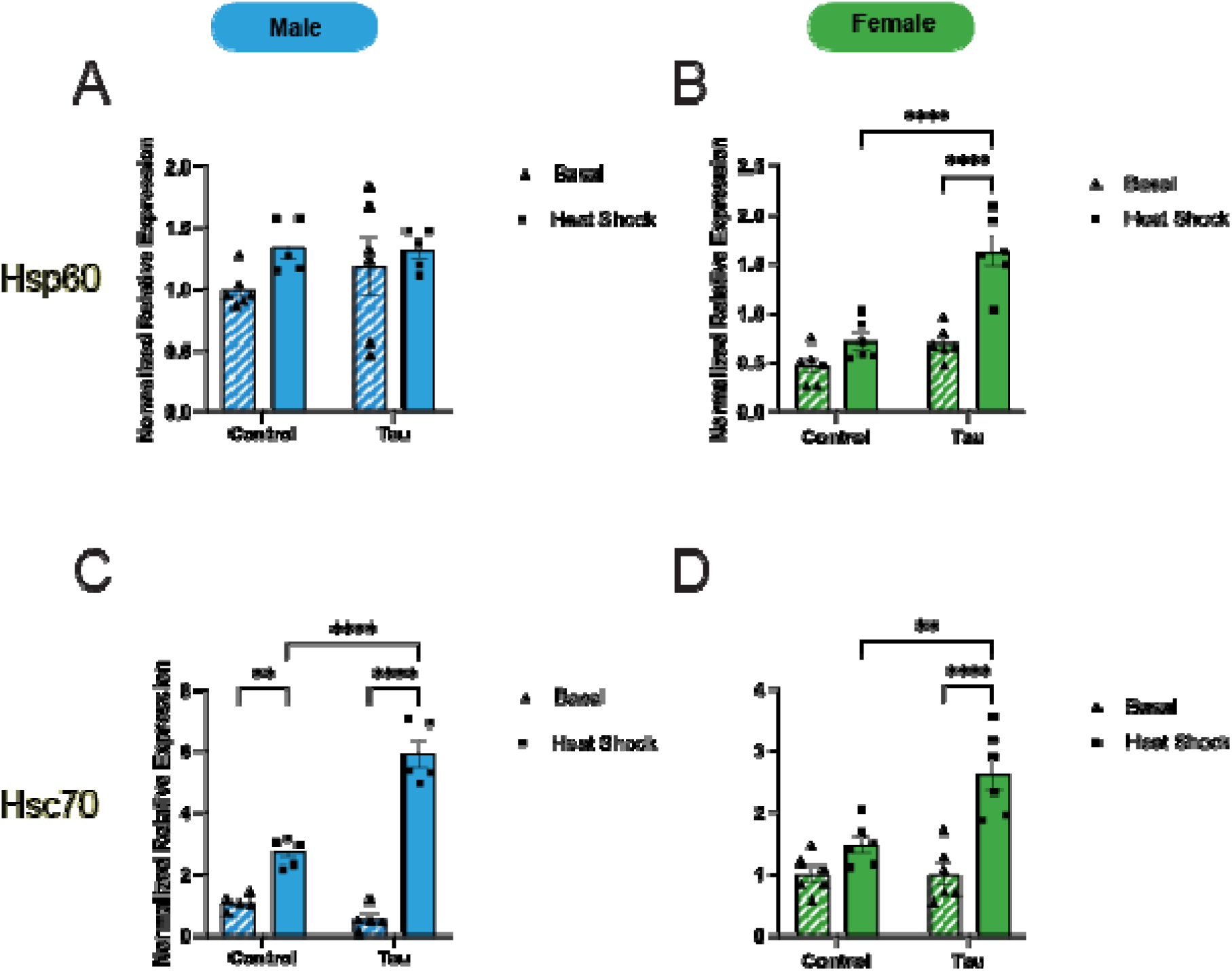
Constitutive Hsp expression changes in male and female flies in response to glial tau expression and heat stress. (A-D) Relative mRNA expression levels of two constitutive heat shock proteins, *Hsp60* (A-B) and *Hsc70* (C-D) for male and female flies under basal and heat stress conditions, comparing control and glial tau transgenic flies. Expression values for are normalized to day 10 basal male control flies for each Hsp. Data are presented as mean + SEM (n = 6) and analyzed using two-way ANOVA with Tukey’s multiple comparisons, * p < 0.05; ** p < 0.01; *** p < 0.001, **** P < 0.0001.

*Hsc70* expression increased significantly response to heat stress in control male flies, but not females, heat stress induced *Hsc70* expression in both sexes of tau transgenic flies (Figure 4C, D). The combined presence of heat stress and glial tau expression further enhanced *Hsc70* expression in both males and females, suggesting an additive effect of these stressors on *Hsc70* regulation (Figures 4C, 4D).

### Differential Hsp protein expression is observed in response to heat stress and glial tau expression

To determine if the observed changes in RNA expression corresponded with changes in protein expression, we examined Hsp27 and Hsp70 protein levels by western blot analysis. Protein expression was compared across sex, heat stress, and glial tau expression conditions. Quantified protein levels were normalized to male basal conditions. Under basal conditions, Hsp27 protein levels remained unchanged in both males or females (Figure 5A, B). Heat stress induced an ∼3-fold increase in Hsp27 expression in both sexes, regardless of glial tau expression (Figure 5A, B). However, a sex-specific difference emerged under heat stress in the presence of glial tau where females expressing glial tau exhibited significantly higher Hsp27 levels than males under the same conditions (Figure 5A, B).

**Figure 5.**
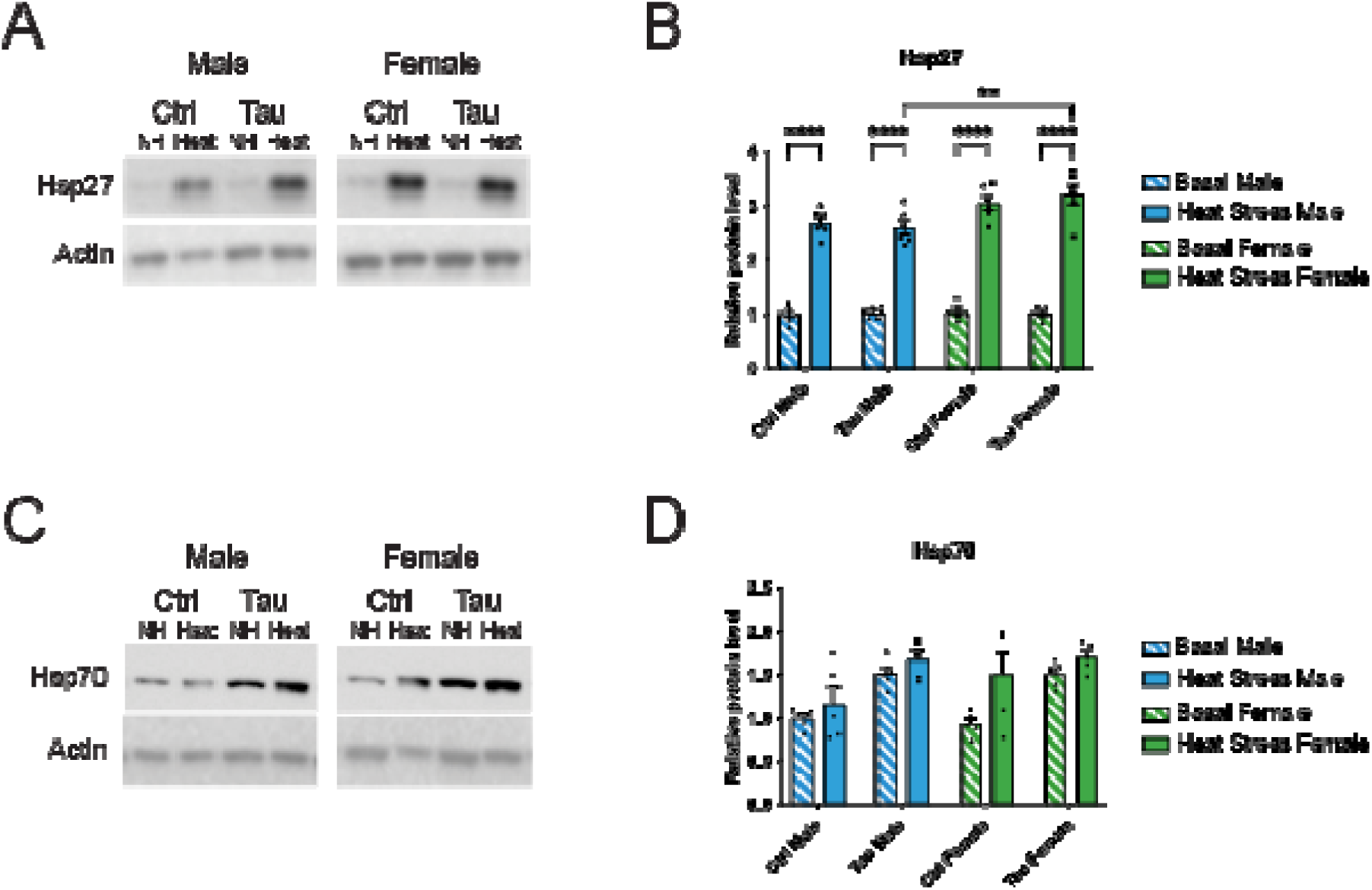
Hsp27, but not Hsp70, protein levels are increased in response to heat stress in a sex-dependent fashion. (A) Representative western blot images of Hsp27 levels and actin levels from brain lysates of male and female control and tau transgenic flies in response to heat stress. (B) Quantification of Hsp27 levels by densitometric analysis, expressed as a ratio of Hsp27 to actin. (C) Representative western blot images of Hsp70 levels and actin levels from brain lysates of male and female control and tau transgenic flies in response to heat stress. (D) Quantification of Hsp70 levels by densitometric analysis, expressed as a ratio of Hsp70 to actin. Protein levels are normalized to day 10 basal control male flies for each Hsp. Data are presented as mean + SEM (n = 5 - 6) and analyzed using two-way ANOVA with Tukey’s multiple comparisons, * p < 0.05; ** p < 0.01; *** p < 0.001, **** P < 0.0001.

In contrast, Hsp70 protein expression was more variable than Hsp27 and did not show a consistent or significant response to heat stress or glial tau expression across sexes (Figure 5C D). While western blot analysis of individual flies suggested a possible increase in Hsp70 levels with glial tau expression in males (Figure 5C), this trend was not statistically significant when averaged across all biological samples (Figure 5D). Together, these results indicate that Hsp27 protein expression reflects both heat stress and glial tau-related regulation in a sex-dependent manner, while Hsp70 protein expression appears more variable and less responsive to these conditions.

## Discussion

The Hsp chaperone network promotes proteostasis by fostering protein-protein interactions among member and client proteins. (12, 45, 46) Dysregulation of this network contributes to the development of tauopathies and other protein-misfolding diseases, although the mechanisms underlying this process remain unclear. (47, 48) Furthermore, many neurodegenerative diseases exhibit distinct sexual dimorphisms in incidence and progression.(39, 49, 50) In this work, we examined how sex, heat stress, and early glial tau expression affect inducible (*Hsp22*, *Hsp23*, *Hsp26*, *Hsp27*, *Hsp70*, and *Hsp83*) and constitutive (*Hsp60* and *Hsc70*) Hsp expression in the *Drosophila* brain. We observed noticeable differences in Hsp expression patterns across conditions and we found that, generally, females exhibit a more robust chaperone response in response to stress. These results provide the first systematic determination of Hsp expression patterns in response to heat stress and glial tau expression, and provide insight into how stress-responsive pathways may contribute to sex differences in tauopathy progression.(49, 51–53)

### Basal Hsp expression is influenced by sex and glial tau expression

Under basal conditions, in the absence of heat stress, sex differences in Hsp expression were evident (Figure 1). In control flies, males exhibited significantly higher expression of a subset of Hsps (*Hsp23, Hsp26*, and *Hsp70*), while females expressed higher levels of *Hsp27*. In glial tau transgenic flies, males displayed enhanced expression of *Hsp23* and *Hsp27*, in contrast to the elevated levels of *Hsp27* in female control flies (Figure 1). This relative discrepancy in Hsp27 expression in female tau transgenic flies is notable as Hsp27 is a small heat shock protein that functions as a holdase for misfolded or aggregated proteins, and serves as the first line of defense in proteostasis. (54–59). The blunted expression of Hsp27 in female glial tau transgenic flies may reflect a sex-specific vulnerability to tau-induced proteostatic stress. Hsp27 is capable of forming a stable complex with tau, and then recruiting the Hsp70/Hsp40/Hsp90 refolding complex to prevent aggregation. (24, 59, 60) Analogous sex-specific patterns in Hsp27 expression have been reported in other cell types, including mammalian cardiac muscle and blood cells.(61–63). Constitutive Hsps (*Hsp60* and *Hsc70*) did not display sex-specific basal expression patterns in control and glial tau transgenic flies (Figure 1).

### Inducible Hsps are Differentially Regulated by Heat Stress and Tau

As expected, heat stress robustly induced all small and large inducible heat shock proteins in both sexes in control and glial tau transgenic flies. This upregulation was not surprising, however, females displayed greater induction, with ∼2-4-fold higher expression of *Hsp22*, *Hsp26*, *Hsp27*, and *Hsp70* relative to males. Specific instances of sex differences in Hsp expression have been reported, with males displaying elevated Hsp72 expression after exercise (63) and differences due to oxidative stress and aging have been monitored.(64) Further studies have noted a female advantage in stress response (52, 65) characterized by faster adaptation and greater sensitivity to some stress conditions.(66). This has previously been linked directly and indirectly to hormone regulation, including estrogen.(64, 67) Whether or not similar hormone-related effects underlie how sex determines Hsp expression in our studies remains to be determined.

Heat shock proteins are induced in response to a variety of stress conditions including: heat stress, hypoxia, oxidative, and exposure to toxins. We suggest that the presence of misfolded proteins in excess, such as expression of glial tau, are another form of stress on the proteostasis system. Since heat shock proteins serve a protective role during heat stress (14, 68) in addition to a regulatory role during normal cell growth, we further examined the global induction of Hsps during heat stress by sex and +/− glial tau expression. While heat shocked male and female flies exhibited some similarities, striking differences emerged for both small (Hsp22, Hsp23, Hsp26, and Hsp27) and large Hsps (Hsp70 and Hsp83).

Our data indicates the expression of glial tau impacts both Hsp mRNA and protein expression, for a subset of the chaperone network. Male flies exhibit a decrease in *Hsp23* and *Hsp26* in response to heat and glial tau, while female flies exhibit an increase in *Hsp27*, *Hsp70*, *Hsp83*, *Hsp60*, and *Hsc70* as a result of glial tau expression in the presence of heat stress. There is no observed effect for Hsp27 or Hsp70 (Figure 5) due to tau expression. We hypothesize that the effects observed at the mRNA level would be observed at the protein level as well, but at a delayed timepoint.

Tau is a known client protein for several Hsps.(32, 56, 58, 60, 69–71) For example, Hsp70 binds to tau at a location that overlaps with a portion of the microtubule-binding site on tau, (72) suggesting that Hsp70 binding may regulate physiological tau function and prevent aggregation. Our results show that the presence of tau in day 10 flies augments Hsp70 expression in females. The elevated levels of Hsp70 may have implications in Hsp70-mediated control of tau aggregation. Moreover, Hsp70 levels play a critical role in general protein folding, and the higher Hsp70 levels observed in females may impair general hydrophobic collapse (73, 74). Therapeutic strategies involving inhibition of Hsp70 have been evaluated to facilitate tau clearance (24, 69)and our results will enable more refined strategies based on sexually dimorphic expression patterns.

Sex-dependent stress responses were observed for some chaperones, including Hsp27. Females generally expressed Hsp27 more robustly. Hsp27 contributes to the initial response to tau in females, as it has been previously indicated to bind to tau and prevent aggregation.(32, 50, 58, 59) The lack of an increase in male flies in response to tau expression is unexpected, although this may partially explain the previously reported differences in tau load and disease progression which are observed, and distinguish pathology between sexes. (49) Hsp27 is secreted from astrocytes to promote neuroprotection.(12) Our finding that glial tau expression decreases basal Hsp27 levels provides a potential mechanism by which glial tau pathology could contribute to tauopathy pathogenesis and progression. However, we see this only in females, suggesting that the sex-specific decrease in Hsp27 may be pathologically relevant. (33)

Hsp83 is interesting in another regard as it is the Drosophila homolog to Hsp90, which is involved in hormone receptor activation (75), and our results indicate that Hsp83 is significantly upregulated during heat stress and glial tau expression, only in females. This suggests that the impacts of tau expression, stress, and/or sex on Hsp83 regulation may occur upstream of transcription and may be susceptible to a feedback process. We hypothesize that Hsp23 and Hsp26 may be involved in a similar feedback process, but one that only occurs in male flies, further suggesting that early (day 10) tau-chaperone interactions vary across sex. For example, Hsp22 is shown to improve cognition by clearing accumulated tau, suggesting a role for sHsps during early tau expression.(76) Our data indicate tau-chaperone interactions are more nuanced than previously determined at a sex-specific level.

Overall, what is most striking is the lack of concerted Hsp expression in either males or females, which indicates that individual Hsps are precisely regulated in response to stress variables, consistent with reports of dysregulation of Hsps in disease models. (77) These results demonstrate the advantage of comparing multiple Hsps under different conditions, illustrating the lack of redundancy in the chaperone network and suggesting each Hsp is individually transcriptionally regulated. Our results distinguish key sexual dimorphisms in Hsp expression, suggesting that males exhibit a more modest heat shock protein response to heat stress and tau expression compared to females. This may be due to a higher stress threshold required for Hsp activation in males compared to females. Generally, female flies present a more robust heat shock protein response to heat stress and tau expression, indicating the threshold required for activation of the heat shock protein response in females is differentially regulated compared to males. Overall, these results contribute to our understanding of the molecular mechanisms that mediate sex differences in neurological stress response, tauopathies, and aging-related disease.

Hsp27 and Hsp70 preferentially associate with phosphorylated tau under stress conditions and co-immunoprecipitated with tau from AD brain homogenates. (78) Although low-levels of Hsp27 and Hsp70 are present in neurons and glia, glia exhibit higher levels of Hsp27 and Hsp70 in astrocytes and microglia, respectively, relative to neurons. (79) Although there is a robust induction and recruitment of the Hsp27/Hsp70 pathway in glia, this machinery is ultimately insufficient to prevent tau pathogenesis. (80) Some reports suggest that prolonged activation of the Hsp27/Hsp70 chaperone pathway may induce proteotoxicity, while other reports indicate a protective effect provided through expression/induction of Hsp27 (24, 30, 69, 81). Further evidence for Hsp27 neuroprotection suggests the mechanism may be tied to the interactions between Hsp27, tau, and microtubules (47, 72, 82). In male flies, we observe no change in Hsp27/Hsp70 expression under basal conditions, whereas females exhibit a decline in Hsp27 expression as a result of tau expression.

Glial cells appear to participate in the propagation and spread of tau pathology from cell to cell and across brain regions, which may accelerate neurodegeneration. (83–86) Basal expression of *Hsp23* and *Hsp27* in only females is reduced due to tau expression (Figure 1). The relative decrease in female flies may suggest a mechanism by which tau expression decreases the Hsp response, through a sexually dimorphic mechanism. Hsp27 is further known to interact with tau and has previously been shown to rescue neural deficits due to tau expression, but it is not the only chaperone known to interact with tau. (59, 87) This response at day 10 in females, but not males, may indicate different thresholds of a particular “stress” or client protein may be required to initiate a chaperone response. In a different model of stress pathology in mice, alcoholic liver injury induced sexually dimorphic changes in Hsp27 expression, suggesting that sex-related variables or hormones can trigger sexually dimorphic stress-induced expression patterns. (88) The presence of tau during early aging differentially impacts male and female basal heat shock protein expression. Whether this phenomenon directly contributes to variations in disease pathologies remain unknown.

## Conclusions

Our findings reveal an intricate and sexually dimorphic regulation of the heat shock protein response in the aging Drosophila brain, with profound implications for neurodegenerative disease. While female flies exhibit a more robust chaperone response, particularly through Hsp27 and Hsp70 compared to males, this protection is further elevated under the combined burden of stress and tau expression and observed in constitutive Hsp60 and Hsc70. The elevation of Hsp27 and Hsp70 in females expressing tau is particularly notable, as these chaperones are crucial for preventing tau aggregation. The dampened and attenuated response of Hsp23 and Hsp26 expression in males suggests heightened vulnerability to proteotoxic stress. These sexually dimorphic responses could underlie well-documented differences in neurodegenerative disease susceptibility, emphasizing the need for sex-specific therapeutic strategies that account for the complex interplay between stress and tau pathology.

A breakdown of proteostasis is seen in human tauopathies (89–93), indicating the early response observed in females may contribute to an overburdened proteostasis network that is impacted during aging. Future studies will evaluate the impacts of aging on this system and alterations in mRNA and protein expression due to aging, stress, and sex. Further downstream processes, specifically the ubiquitin-proteasome pathway, which demonstrates a decline in function with age may also be evaluated. Moreover, proteins are prone to age-related damage including: cleavage, covalent modifications, oxidative lesions, glycation, cross-linking, and denaturation. As the cell ages, mitochondrial malfunction and the resulting decrease in ATP production and increase in reactive oxygen species (ROS) can lead to a greater output of misfolded proteins and more aggregation, which impacts the broader proteostasis network. Therefore, studies that characterize the impacts of aging, stress, and tau on heat shock proteins will enable a more detailed understanding of disease-related pathologies.

## Materials and methods

### Drosophila melanogaster stocks and genetics

*w^1118^* was obtained from the Bloomington Stock Center (BL#3605), and double recombinant control flies (*repo-GAL4, tubulin-GAL80^TS^/TM3,Sb*) and triple recombinant tau flies (*repo-GAL4, tubulin-GAL80^TS^, UAS-Tau^WT^/ TM6B,Tb*) were used (Scarpelli et al., 2019). Flies were maintained at 25°C in plastic vials with standard cornmeal-based food (NutriFly) supplemented with propionic acid and tegosept. Crosses (*repo-GAL4, tubulin-GAL80^TS^/TM3,Sb* X *w^1118^* and *repo-GAL4, tubulin-GAL80^TS^, UAS-Tau^WT^/ TM6B,Tb* X *w^1118^*) were performed at 25°C in incubators with humidity control and 12-hour light:dark cycles. Progeny were selected against balancers and separated by sex, and aged to 10 days at 25°C. Flies were flipped to new food every 2-3 days while aging.

#### *Drosophila* Heat Stress Conditions

Once flies were aged to ten days, half of the vials were kept at 25°C (no heat (NH), basal conditions), while the other half were placed at 37°C for 1 hour to induce heat stress (HS). Following the heat stress period, flies were placed back to 25°C for a 30 min recovery period and processed for RNA extraction.

#### RNA Extraction and cDNA Synthesis

Ten day old basal and heat stress control and tau transgenic flies were anesthetized with CO_2_ and flash-frozen in a dry ice/ethanol bath. Heads were dissected and homogenized in TRIzol. Each biological replicate contained 5-10 fly heads. RNA was extracted from these samples via chloroform/isopropanol (Life Technologies) RNA precipitation, according to standard protocols.(94) Briefly, 0.2 volumes of chloroform were added to each TRIzol homogenate, followed by phase separation in Phase Lock Gel-Heavy 2mL tubes after spinning at 4°C, 12k rpm for 1 min. Isopropanol was added and the solution was precipitated overnight at −20°C and subsequently spun at 4°C, 13.6k rpm for 30 min. The pellet was washed with 800 μL of 75% ethanol and spun for 10 min at 13.6k rpm. The pellet was redissolved in nuclease-free H2O and placed in a 60°C heat block for 10 min to facilitate dissolution. RNA concentration and purity were analyzed using a NanoDrop 1000 spectrophotometer, and RNA was ≥ 40 ng/μL. Samples were treated with DNase (DNA-free kit; Ambion) and 250 ng of RNA was used to generate cDNA using the SuperScript First-Strand Synthesis System kit (Invitrogen).

#### Quantification of Hsp expression by qPCR

qPCR was performed using SYBR Green (Applied Biosystems). Each condition represents the data from 5-6 biological replicates, each composed of three technical replicates. *RpL32* was used as the internal control gene, and primer efficiency for all primer pairs was determined to be between 87-108%. Primers used were: *Hsp22*: (Forward) *GAT GAA CTG GAC AAG GCT CTA A*, (Reverse) *TAT GAT TGG CGA CTG CTT CTC*; *Hsp23*: (Forward) *GCG ATA ACA GCT AAA GCG AAA G,* (Reverse) *CAA GGC TCA ACA ATG GAA TA*; *Hsp26*: (Forward) *TGG ACG ACT CCA TCT T,* (Reverse) *TAG CCA TCG GGA ACC TTG TA*; *Hsp27*: (Forward) *GAA GTC GTG AAG GAG GAA G,* (Reverse) *GGC AAC ACT CCC GTT TCT*; *Hsp60*: (Forward) *AGA TGT GAT GAG AAC CGA AAC C*, (Reverse) *CCG ACT GCT GAT GAC TGA TAA C*; *Hsp70A*: (Forward) *GTC GTT ACC GAG GAA C*, (Reverse) *CAC CTT GCC ATG TTG GTA GA*; *Hsc70*: (Forward) *CCT ATG TTG CCT TCA CCG ATA C*, (Reverse) *TCG AAC TTG CGA CCA ATC AA*; *Hsp83*: (Forward) *CAC ATG GAG GTC GAT TAA G*, (Reverse) *CGG CCG TAG TAA ACT CAG TAT AAA*; *RpL32*: (Forward) *CCA GTC GGA TCG ATA TGC TAA G*, (Reverse) *CCG ATG TTG GGC ATC AGA TA*

#### Western Blotting and Protein Quantification

Western blot analysis was performed to quantify Hsp27 and Hsp70 protein expression across all conditions including sex, genotype, and heat stress. Single heads isolated from flash frozen day 10 flies were homogenized in 2x Laemmli sample buffer (65.8 mM Tris-HCl, pH 6.8, 2.1% SDS, 26.3% (w/v) glycerol, and 1% bromophenol blue), and heat denatured for 10 minutes. Samples were subjected to SDS-PAGE in a 26-well polyacrylamide gel (BioRad), using one head per lane. Protein was transferred to a polyvinylidene fluoride (PVDF) membrane using the Trans-Blot Turbo Transfer System (Bio-Rad) and blocked in 2% milk with 0.1% Tween® in TBS. Membranes were incubated overnight in primary antibodies, washed in 2% milk with 0.1% Tween® in TBS (wash buffer), followed by incubation in a complementary secondary antibody. Secondary antibodies were rinses and ECL substrate (Bio-Rad) was utilized for imaging. The signal was detected by chemiluminescence imaging on an Azure Biosystems c600.

The following primary and secondary antibodies were used: ms ⍰ Actin (1:1,000) DSHB, ms ⍰ Hsp27 (1:1,000) Abcam, ms ⍰ Hsp70 (1:1,000) (gifted by R.Tanguay (95)), gt ⍰ rb IgG(HRP) (1:20,000) ThermoFisher, gt ⍰ ms IgG (HRP) (1:20,000) ThermoFisher. Protein bands in western blot images were quantified with a densitometric analysis in ImageJ. Protein band densities were quantified relative to a corresponding actin loading control, to account for loading variability. Normalized density was achieved by comparing the relative band density to the average band density of all basal control male samples.

#### Statistical Analyses

All statistical analyses were performed in GraphPad Prism 10 software. Relative expression of mRNA is determined by normalizing male or female flies (respectively) to basal conditions, unless otherwise indicated. All data presented are expressed as mean ± SEM. Multiple group comparisons analyzing sex, heat stress, tau expression, and aging variables were evaluated with a Two-Way ANOVA, followed by Tukey’s multiple comparisons tests.

## Abbreviations

AD: Alzheimer’s disease
Hsc: Heat shock cognate
Hsp: Heat shock proteins

## Acknowledgments

We thank members of the Colodner and McMenimen labs for feedback and constructive discussions.

## Author Contributions

Margot Whitmore: data curation, formal analysis, investigation, methodology, writing-review and editing, Maeve Coughlan: data curation, formal analysis, investigation, methodology, Martha A. Kahlson: data curation, investigation, methodology, writing-review and editing, validation, Jaasiel Alvarez: data curation, formal analysis, investigation, methodology, Louisa Zebrowski: data curation, investigation, Kathryn A. McMenimen: Conceptualization, Data curation, Formal analysis, Funding acquisition, Investigation, Methodology, Project administration, Resources, Supervision, Validation, Visualization, Roles/Writing - original draft; and Writing - review & editing, Kenneth J. Colodner: Conceptualization, Data curation, Formal analysis, Funding acquisition, Investigation, Methodology, Project administration, Resources, Supervision, Validation, Visualization, Roles/Writing - original draft; and Writing - review & editing

Declaration of Competing Interests: The authors declare no competing financial interests.

